# White spot syndrome virus infection induces Caspase 1-mediated antiviral cell death in crustacean

**DOI:** 10.1101/322982

**Authors:** Geng Yang, Jiajia Wang, Tao Luo, Jun Wei, Yadong Yang, Wenyuan Zhang, Xiaobo Zhang

## Abstract

In vertebrates, pyroptosis is an intensely inflammatory form of programmed cell death which is dependent on Caspase 1 activation and release of cytoplasmic cytokines including IL-1β. This death pathway is critical for controlling pathogenic infection by mobilizing immune cells and stimulating the development of adaptive immune response. In invertebrates, however, due to the lack of adaptive immune response, it is still elusive whether Caspase 1-dependent cell death pathway exists. In this study, our data showed that Caspase 1-mediated cell death was activated by white spot syndrome virus (WSSV) infection to control the virus in shrimp. Caspase 1 had a higher expression level in hemocytes and lymphoid-like organ in shrimp and WSSV infection was significantly promoted upon the inhibition of Caspase 1 enzymatic activity. IL-1β-like protein was identified as the substrate of Caspase 1 and its interaction with Caspase 1 was validated ectopically and endogenously. Moreover, IL-1β like protein was released into extracellular contents under WSSV infection and Prophenoloxidase system was activated, resulting in the reduction of WSSV copies *in vivo*. Our data unraveled a previously unidentified mechanism through which Caspase 1-dependent cell death controlled virus infection in shrimp. Therefore, our study opened the possibility that an invertebrate cytokine network might be operative and regulate host defenses against virus infection as in vertebrates.

**Author Summary:** The elimination of infected cells by programmed cell death is considered as one of the most important anti-pathogen responses. Pyroptosis is one form of programmed cell death, which is mediated by Caspase 1 activation. Activated Caspase 1 can process the inactive precursors of interleukin 1β (IL-1β) into mature inflammatory cytokines to induce potent immune responses. Meanwhile, pyroptosis also induces morphologic changes, DNA fragmentation and chromatin condensation to destroy infected cells. However, due to the lack of efficient adaptive immune responses, it is still unclear whether Caspase 1-mediated cell death also exits and plays such an important role in defending hosts from pathogen invasion in invertebrates. In our data, Caspase 1 gene was characterized and Caspase 1-mediated cell death pathway was validated in shrimp. The Caspase 1-mediated cell death pathway was enhanced by white spot syndrome virus (WSSV) infection to control the virus in shrimp. IL-1β-like protein was cleaved by Caspase 1 and released into extracellular contents under WSSV infection to activate Prophenoloxidase system, leading to the reduction of WSSV copies in shrimp in vivo. On the other hand, silencing of *Caspase 1* or *IL-1β-like* gene facilitated virus infection in shrimp. Our study indicated that Casapase-1-mediated cell death played an important antiviral role in shrimp against WSSV infection and opened the possibility that an invertebrate cytokine network might be operative.

## Introduction

One of the most important anti-pathogen responses consists of the elimination of the infected cells by programmed cell death, a response found in all metazoans (1, 2). Apoptosis is perhaps the most widely recognized programmed cell death, and is defined by the requirement for particular cysteine-dependent aspartate-specific proteases, commonly Caspases 3/7, which produce an orchestrated disassembly of the cell resulting in mounting more severe immune responses (3, 4). It is found that apoptosis plays a very important role in determining the outcome of host-pathogen interactions (5, 6). Other than Caspase 3/7-mediated apoptosis, pyroptosis is a more recently identified pathway of host cell death that is stimulated by a range of pathogen infections and non-infectious stimuli (7, 8).

Unlike apoptosis, pyroptosis occurs after Caspase 1 activation. Caspase 1 is usually activated by protein complexes termed inflammasomes in mammals (9). Two types of inflammasomes formed by Nod-like receptors (NLRs) have been identified. One type is NLRC4 and murine NLRP1b and the other type is NLRP3 (10). NLRC4 and murine NLRP1b contain CARD domains that directly interact with the Caspase 1 CARD and then activate Caspase 1 proteolytic activity. In contrast, NLRP3 contains a Pyrin signaling domain instead of a CARD domain (11). The Pyrin domain binds the Pyrin domain of the adaptor protein ASC which is composed of only a Pyrin and a CARD domain. ASC then recruits Caspase 1 via CARD-CARD interactions and finally activate Caspase 1 proteolytic activity (12). Activated Caspase 1 can recognize and process the inactive precursors of interleukin 1β (IL-1β) into mature inflammatory cytokines, which is then secreted into intracellular contents (12). IL-1β is an important mediator of the inflammatory response of the host against pathogen infection (13). Meanwhile, pyroptosis also induces morphologic changes, DNA fragmentation and chromatin condensation to destroy infected cells (7, 9).

During pathogen infection, the host benefits from pyroptosis. The compromised cells are eliminated and more severe immune responses are launched, effectively destroying the protective environment where infectious agents can thrive (14). In Caspase-1-deficient macrophages where pyroptosis is compromised, higher intracellular bacterial loads are detected during *L. pneumophila* infection (14). Pyroptosis also promotes pathogen clearance by acting as an alarm signal that recruits immune cells to the site of infection (15). The secretion of IL-1β and IL-18 promotes leukocyte activation and immuno-stimulatory factors are released from lysed cells into the extracellular milieu (15). A bunch of released cytosolic products are potent damage-associated molecular patterns (DAMPs), such as high-mobility group box 1 (HMGB1), heat-shock proteins and DNA-chromatin complexes. By the activation of pattern-recognition receptors, these DAMPs accelerate proinflammatory cytokine production (16). On the other hand, pathogens have evolved effector proteins capable of directly inhibiting Caspase 1 and inflammasome activation, preventing cells from pyroptosis (17, 18). Virulent *Pseudomonas* strains express the effector protein ExoU, which blocks Caspase 1 activation and *Y. pseudotuberculosis* T3SS Rho-GTPase activating protein YopE inhibits inflammasome activation (17, 19). The host-pathogen interaction mechanisms suggest that the activity of Caspase 1 leading to pyroptosis may represent an important antiviral pathway that prevents replication and spread of viruses to neighboring cells.

Pyroptosis is such a potent mechanism to clear intracellular pathogens. However, current studies on the role of pyroptosis in host-virus interactions has not been extensively explored and it is still unclear whether pyroptosis also exits and plays such an important role in defending hosts from pathogen invasion in invertebrates. In this study, we aim to find out whether pyroptosis is also one of the pathways that contributes to eliminating virus in invertebrate. The results revealed that the activity of Caspase 1 in shrimp was enhanced by white spot syndrome virus (WSSV) infection. The silencing of *Caspase 1* or *IL-1β-like* gene facilitated virus infection in shrimp. Our study indicated that Casapase-1-mediated cell death played an important antiviral role in invertebrate.

## Results

### Involvement of Caspase 1 in the virus infection to shrimp

In order to investigate whether Caspase 1-mediated cell death plays a role in the immune response of invertebrates to virus infection, the profiles of *Caspase 1* gene was characterized in the course of shrimp immunity against WSSV infection. The sequence analysis revealed that the shrimp Caspase 1 contained caspase activity domains (Fig 1A). The expression of the gene in different organs was then examined by real-time PCR. The result showed that Caspase 1 expression was most significant in hemocytes followed by lymphoid-like-organ, however, we can barely detect the mRNA of Caspase 1 in muscle and heart (Figure 1B). The expression pattern of the gene was further confirmed by detection of protein levels as shown in Figure 1C. These results indicated the shrimp Caspase 1 might function in immunity system--hemocytes and lymphoid-like organ in shrimp, where fighting off invading pathogens usually took place, so the result indicated that Caspase 1 might play a role in the antiviral cellular pathway..

To explore the role of Caspase 1 in virus infection to shrimp, the expression of Caspase 1 was detected at different time points after shrimp were infected with WSSV. The results revealed that the transcriptional level of Caspase 1 was significantly upregulated with the prolonged infection time (Fig 1D). Meanwhile, the protein level of the gene also showed similar expression patterns during WSSV infection (Figure 1E).Taken together, these results indicate that Caspase-1-mediated cell death is involved in virus infection of shrimp.

**Fig 1.**
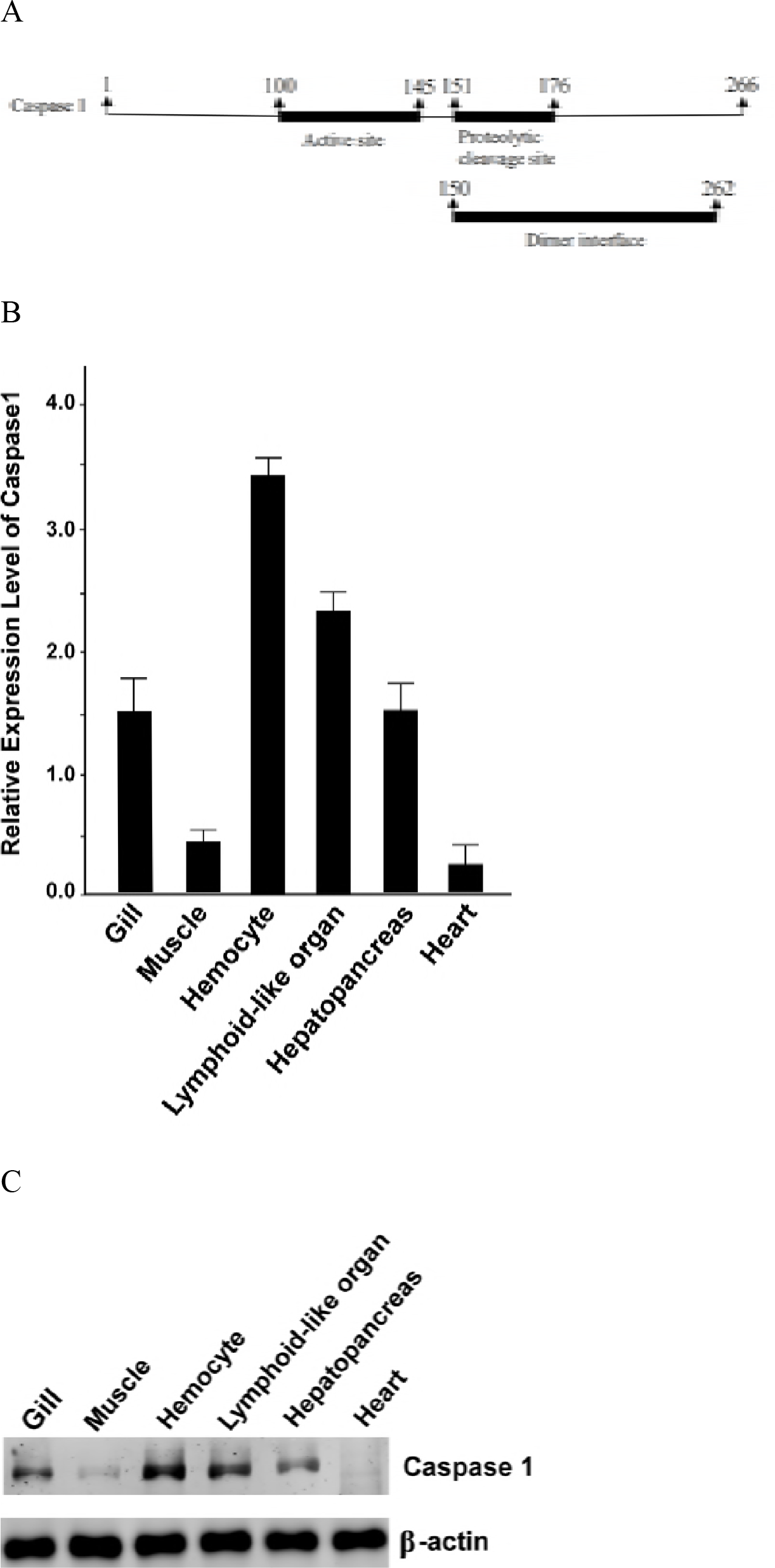
Involvement of Caspase 1 in the virus infection to shrimp. (A) Sequence analysis of shrimp Caspase 1. *Casapse 1* gene was cloned from shrimp hemocytes. Shrimp Caspase 1 contained three domains including active site, proteolytic cleavage site and dimer interface. (B) Detection of Caspase 1 mRNA in different tissues of shrimp. The Caspase 1 mRNA was examined by quantitative real-time PCR. Shrimp β-actin was used as an internal control. (C) Examination of Caspase 1 protein in different shrimp tissues. The Caspase 1 protein was detected by Western blot. Shrimp β-actin was used as an internal control. (D) Expression level of Caspase 1 mRNA in shrimp challenged with virus. Shrimp were challenged with WSSV. At different time post-infection, the Caspase 1 mRNA of shrimp hemocytes were analyzed by quantitative real-time PCR (*, *p*<0.01). (E) Determination of Caspase 1 protein level in WSSV-infected shrimp. Proteins were extracted from virus-infected shrimp hemocytes and analyzed by Western blot. The numbers indicated the time post-infection. Shrimp β-actin was used as an internal control.

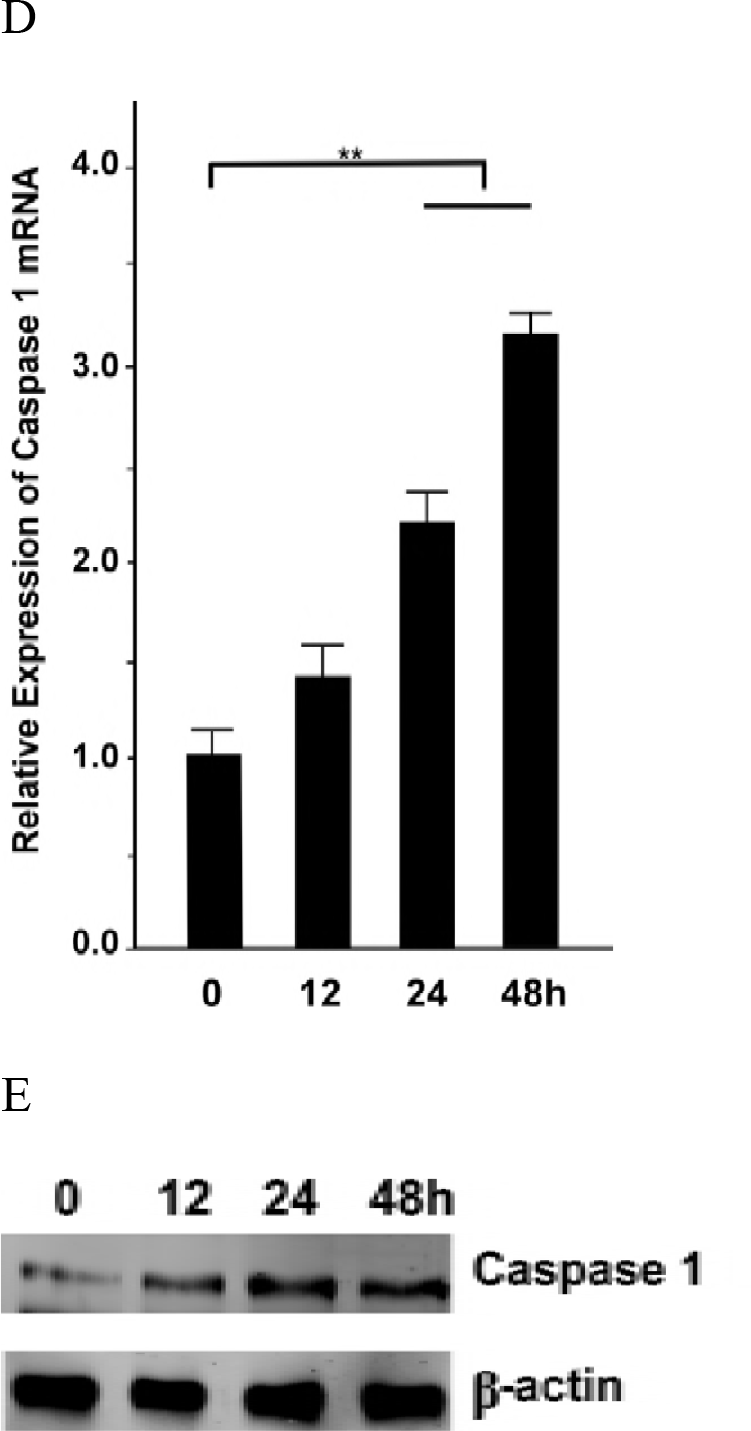

### Requirement of enzymatic activity of Caspase 1 in virus infection

To determine whether the enzymatic activity of Caspase 1 was associated with virus infection, shrimp were challenged with increasing concentrations of WSSV and hemocytes were collected for the detection of Caspase 1 activity by incubating with the substrate Ac-YVAD-pNA. The results showed that Caspase 1 activity was greatly enhanced while more viruses were used for infection (Fig 2A). Moreover, Caspase 1 activity increased dramatically along with the prolonged infection time (Fig 2B). These results indicated that the enzymatic activity of Caspase 1 was actually indispensible in the pathway of virus infection.

**Fig 2.**
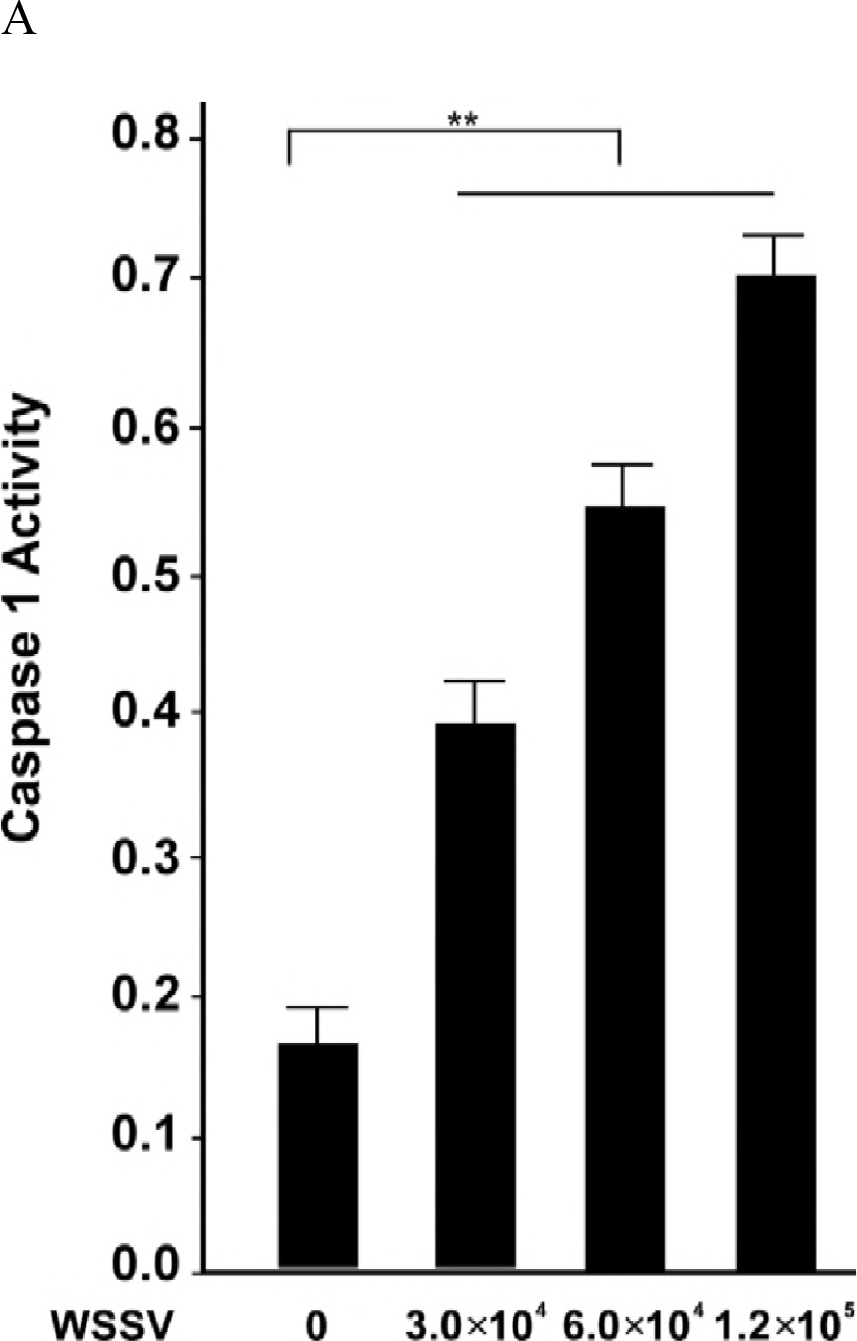
Requirement of enzymatic activity of Caspase 1 in virus infection. (A) Influence of WSSV concentration on Caspase 1 activity. Caspase 1 activity was measured at OD_450_ after shrimp were challenged with increasing WSSV concentration. Caspase 1 activity was detected at 48 h post-infection. (B) Effects of virus infection on Caspase 1 activity. Caspase 1 activity was measured at different time points after shrimp were challenged with WSSV. (C) Impact of inhibition of Caspase 1 activity on virus infection. WSSV copy number was measured by quantitative real-time PCR at 48 h after shrimp were treated with Caspase 1 inhibitor with or without WSSV infection.

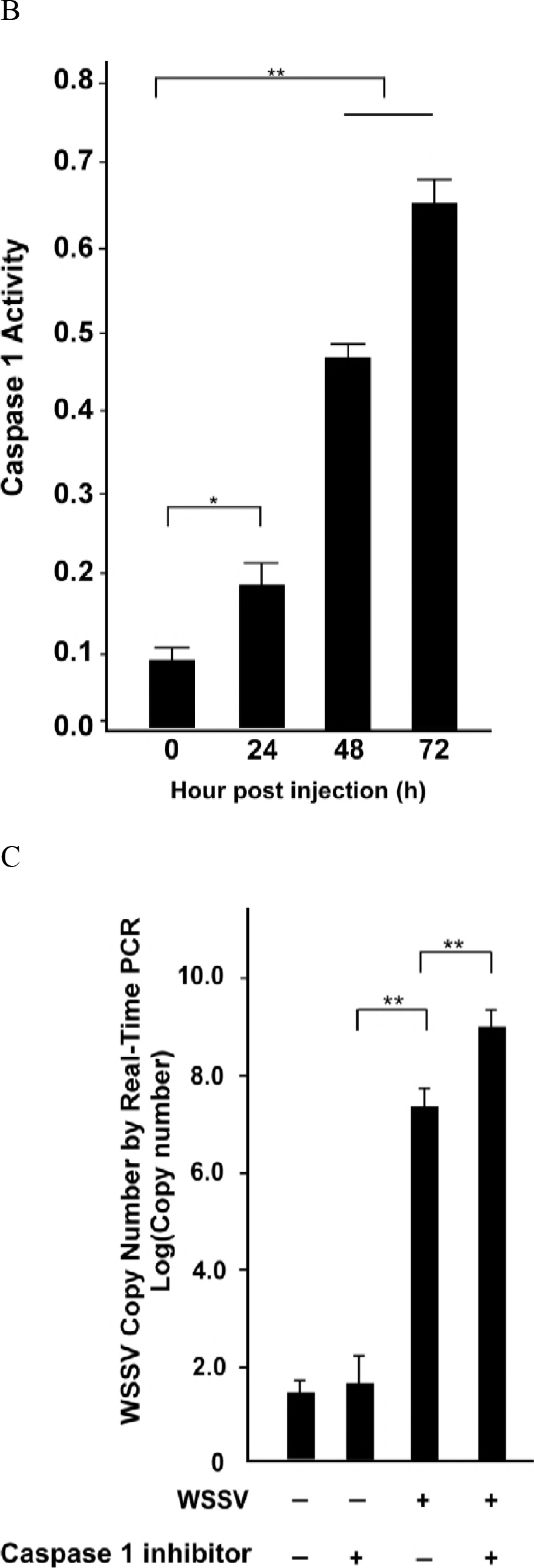

To explore whether the Caspase 1 activity facilitated or inhibited virus infection, Caspase 1 inhibitor Belnacasan was used in combination with WSSV injection or not, and WSSV copy number was then measured by quantitative real-time PCR. The data showed that the WSSV copy number increased significantly upon the inhibition of Caspase 1 enzymatic activity, suggesting that Caspase 1 played an antiviral role possibly conducted by its enzymatic activity.

### Interaction between Caspase 1 and IL-1β like protein

In order to reveal the cellular pathway in which Caspase 1 participated to play an antiviral role, we performed Coimmunoprecipitation (Co-IP) to identify the substrate of Caspase 1. The Co-IP data showed that one specific band was discovered after Caspase 1-specific antibody but not IgG was used for CoIP, which was then identified as IL-1β like protein by mass spectrometry (Fig 3A). To further demonstrated the interaction between Caspase 1 and IL-1β like protein, Flag-Caspase 1 and His-IL-1β like protein was respectively cloned into PIZ/V5-His plasmid and co-transfected into insect High Five cells. Anti-Flag IP was performed and then proteins were analyzed by Western blotting. The result indicated that His-IL like protein was actually detected (Fig 3B), showing the interaction between Caspase 1 and IL-1β like protein. Meanwhile, anti-His IP was also performed and Flag-Caspase 1 could also be found by Western blotting (Fig 3C). These results revealed that the two proteins could interact with each other by their expressions in insect cells.

To further confirm the interaction Caspase 1 and IL-1β like proteins *in vivo*, endogenous IP was performed using Caspase 1 or IL-1β like protein antibody. The results showed that IL-1β like protein was detected by Caspase 1 endogenous IP and vice versa (Fig 3D).

**Fig 3.**
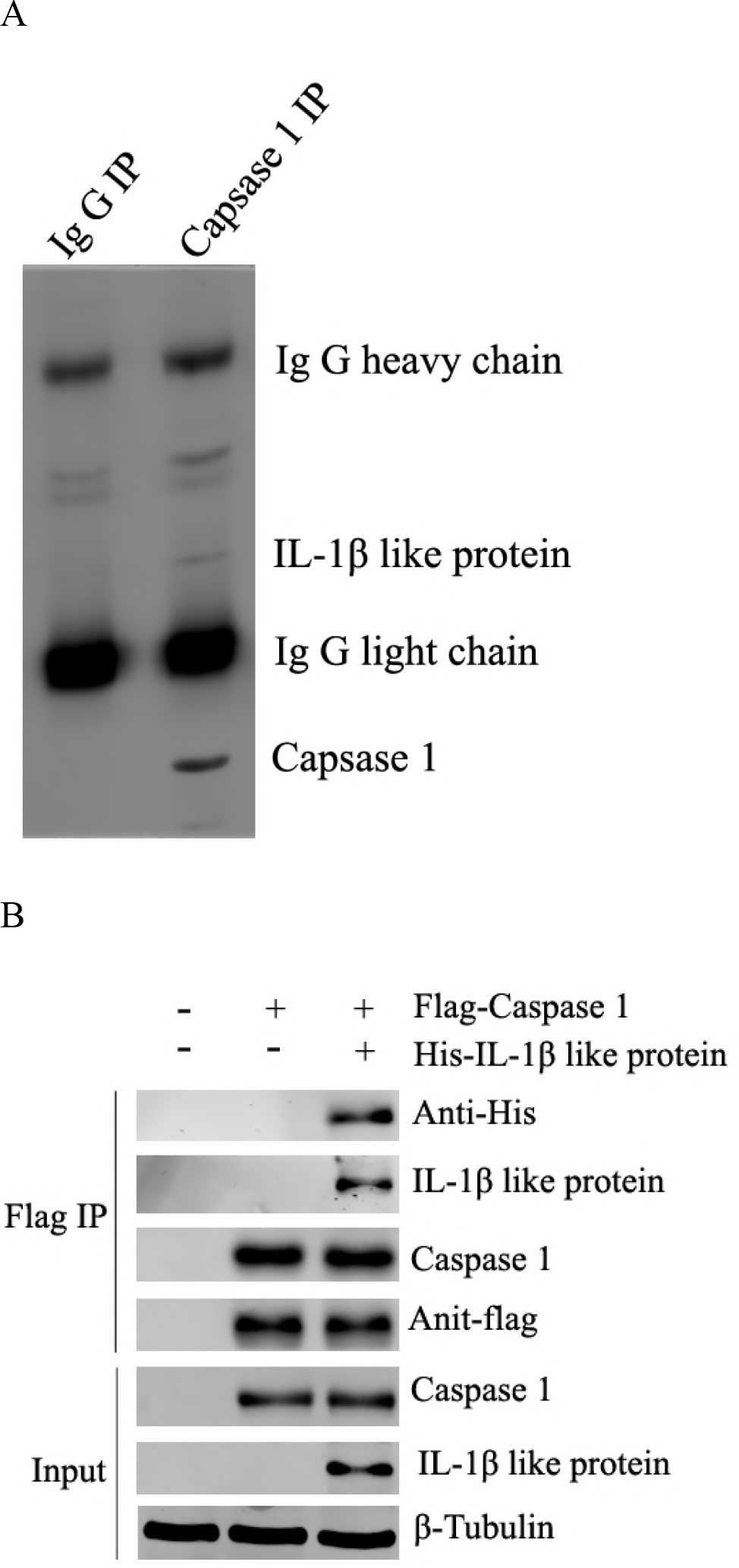
Interaction between Caspase 1 and IL-1β like protein. (A) SDS-PAGE analysis of the elute from Caspase 1 IP by silver staining. Caspase 1 IP was performed and the immunoprecipitate was analyzed by SDS-PAGE by silver staining. The proteins were identified by mass spectrometry. (B) Validation of Flag-Caspase 1 and His-IL-1β like protein by Anti-Flag IP in insect High Five cells. The Flag-Caspase 1 and His-IL-1β like protein constructs were cotransfected into insect cells and Anti-Flag IP was performed. The elute was analyzed by Western blot. β-tubulin was used as an internal control. (C) Validation of Flag-Caspase 1 and His-IL-1β like protein by Anti-His IP in High Five cells. Flag-Caspase 1 and His-IL-1β like protein were cotransfected into High Five cells and Anti-His IP was performed and the elutes was analyzed by western blot. β-tubulin was used as a control. (D) Endogenous Caspase 1 and His-IL-1β like protein interaction by IP. Caspase 1 or IL-1β like protein IP was performed respectively and the immunoprecipitates were analyzed by Western blot. β-tubulin was used as an internal control.

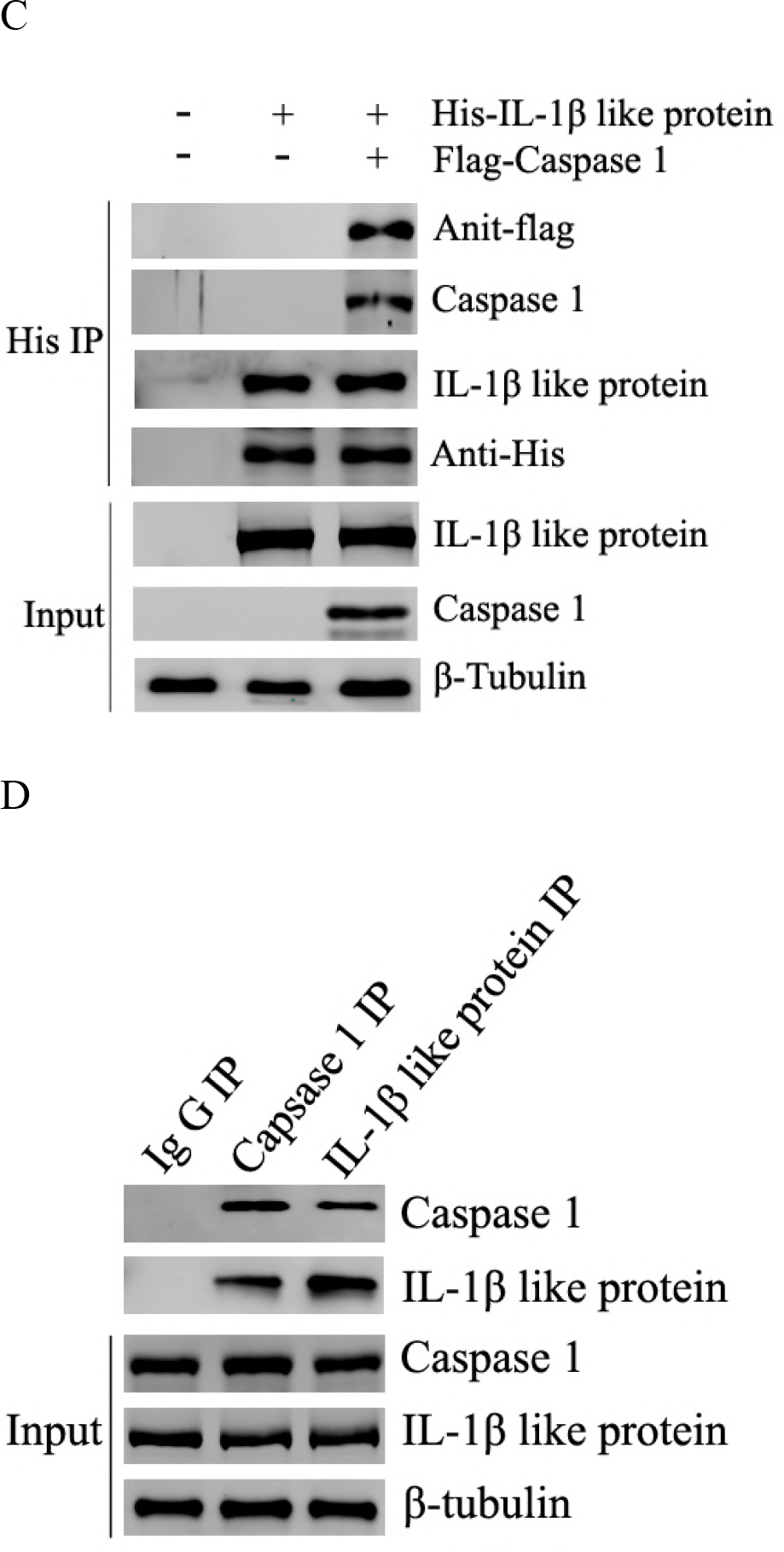

Taken together, these findings identify IL-1β like protein is the substrate of Caspase 1 and the two proteins could interact with each other.

### Redistribution of IL-1β like protein induced by WSSV infection to activate phenoloxidase system

To investigate the functions of Caspase 1 and IL-1β like protein interaction *in vivo*, siRNA respectively targeting *Caspase 1* gene and *IL-1β like protein* gene was designed. The effectiveness of siRNAs was examined by quantitative real-time PCR and Western blotting. From the detected mRNA and protein level, it could be concluded that the siRNAs efficiently knocked down the expressions of *Caspase 1* and *IL-1 like protein* genes (Fig 4A).

**Fig 4.**
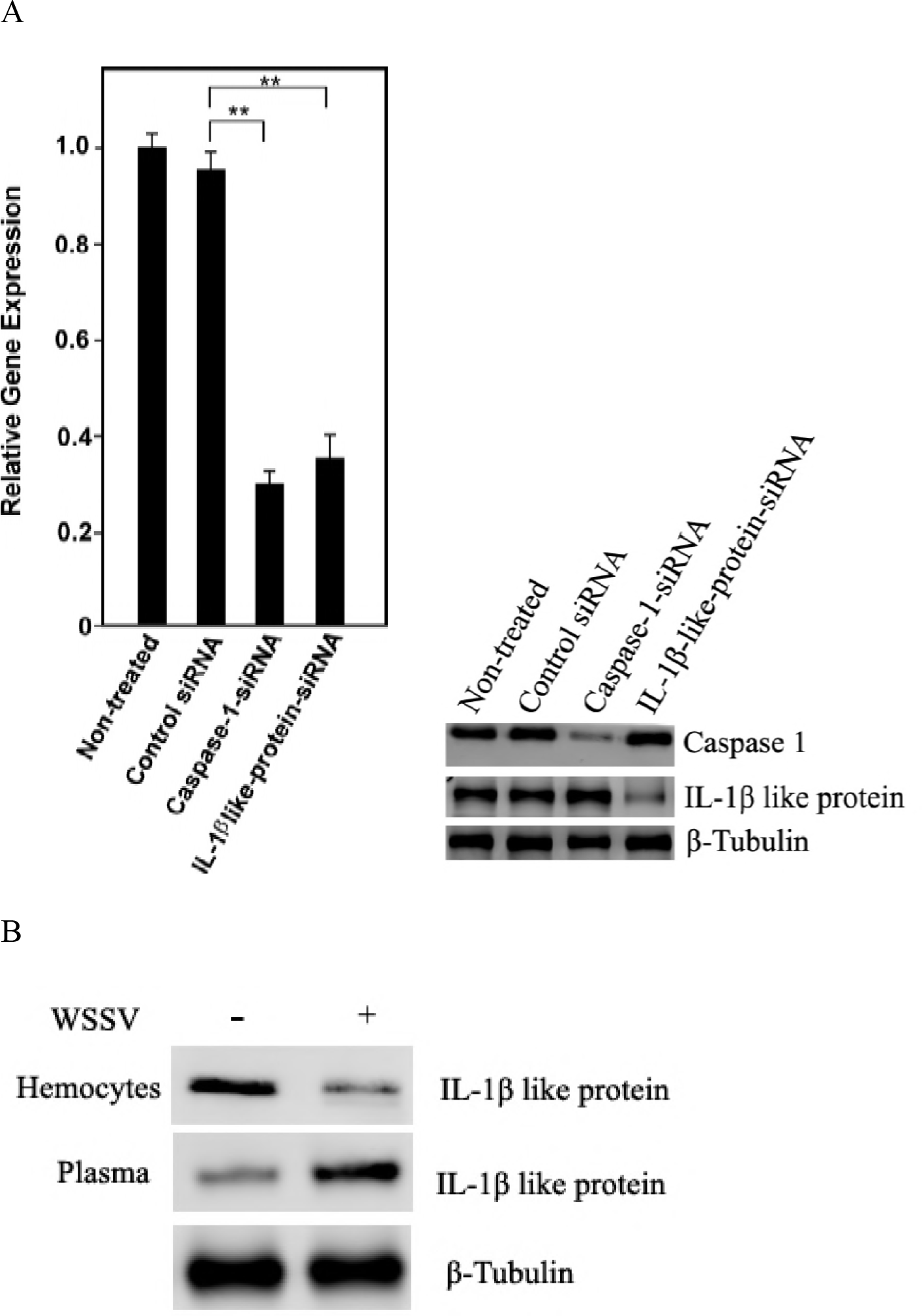
Redistribution of IL-1β like protein induced by WSSV infection to activate phenoloxidase system. (A) Detection of Caspase 1 and IL-1β like protein knockdown efficiency. The mRNA and protein levels of Caspase 1 and IL-1β like protein were respectively detected by quantitative real-time PCR (left) and Western blot (right) at 48 h after shrimp were injected with indicated siRNAs. (B) Detection of IL-1β like protein expression in hemocytes or plasma after WSSV infection. The expression level of IL-1β like protein in hemocytes or plasma was examined by Western blot with or without WSSV infection. Samples were collected at 48 h after WSSV infection. (C) Evaluation of IL-1β like protein expression in hemocytes or plasma after WSSV infection with or without Caspase 1 inhibition. The expression level of IL-1β like protein in hemocytes or plasma was detected by Western blot after WSSV-infected shrimp were treated with Caspase 1 siRNA or inhibitor. Caspase 1 inhibitor or siRNA was immediately injected into shrimp after WSSV infection. Samples were collected 48 h after injection. (D) Examination of phenoloxidase activity upon knockdown of Caspase 1 or IL-1β like protein in shrimp. Phenoloxidase activity was measured at OD_492_ after shrimp were treated with either Caspase-1-siRNA or IL-1β-like-protein-siRNA at 48 h after WSSV injection.

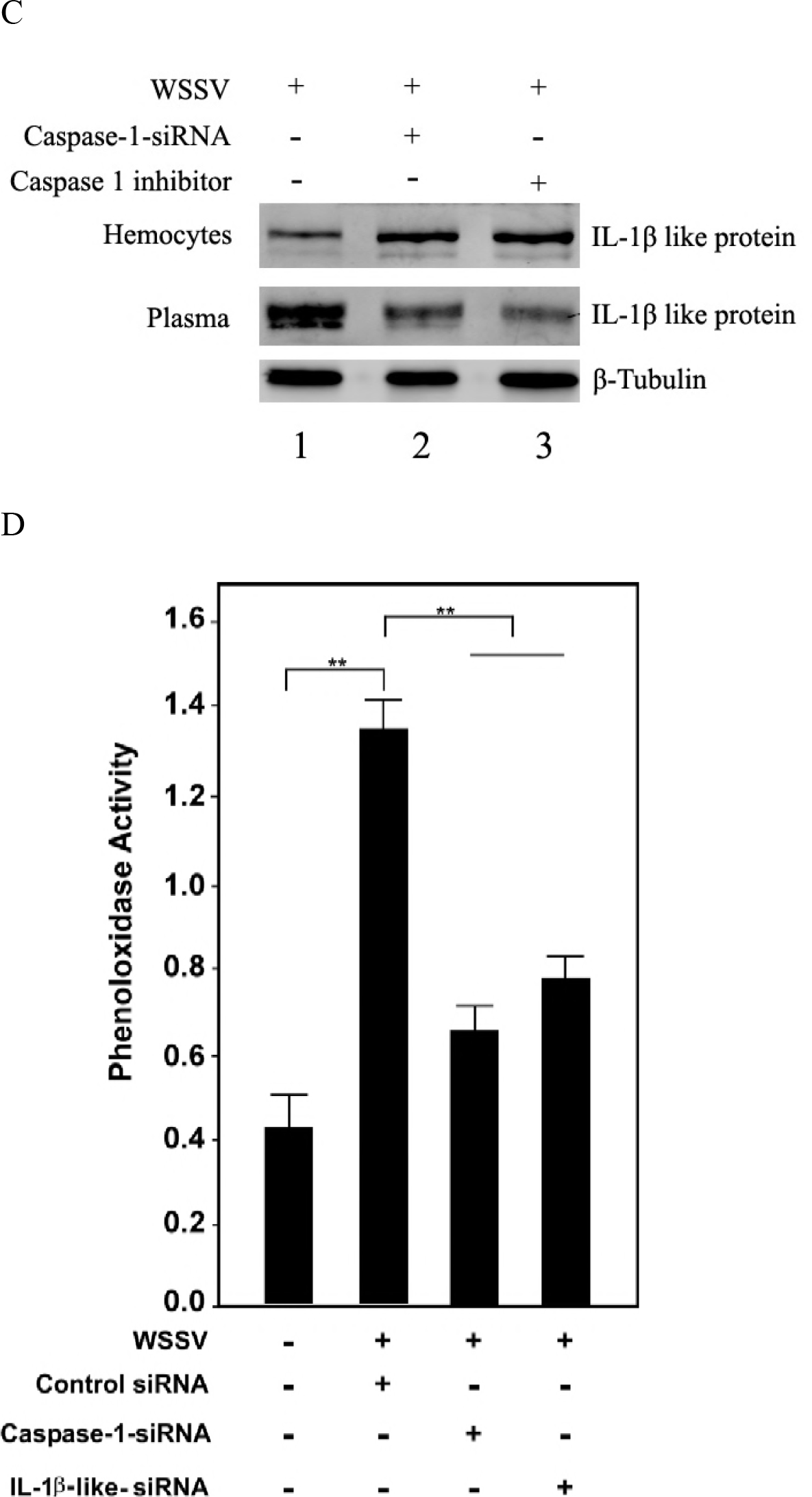

As reported in mammary cells, IL-1β is firstly interacted with and then cleaved by Caspase 1 before it is released into extracellular contents (20), however, in this study, it is still elusive how IL-1β like protein reacts upon virus infection. Therefore, the level of IL-1β like protein in hemocytes and the corresponding plasma before and after WSSV infection was examined. Hemolymph was collected and centrifuged to separate the hymocytes and plasma for the detection of IL-1β like protein. Western blotting analysis revealed that the expression level of IL-1β like protein was increased in the plasma and decreased in hemoctes when shrimp were challenged with WSSV, indicating that IL-1β like protein was released into extracellular contents under virus infection (Fig 4B). In order to determine whether Caspase 1 mediated the release of IL-1 like protein, the expression of Casapse 1 was firstly knocked down by siRNA and then the expression pattern of IL-1β like protein was examined. The results showed that the level of IL-1β like protein in the plasma decreased while the level in the hemocytes increased upon infection with WSSV (Fig 4C). These data showed that the release of IL-1β like protein into plasma was controlled by Caspase 1 in shrimp.

To further determine whether IL-1β like protein was required to be cleaved by Caspase 1 before releasing into extracellular contents, the enzymatic activity of Caspase 1 was inhibited by Belnacasan and then the level of IL-1β like protein was examined. The result showed that the level of IL-1β like protein didn’t increase in the plasma, suggesting that the enzymatic activity of Caspase 1 was required for the release of IL-1β like protein and that IL-1β like protein needed cleavage by Caspase 1 before releasing into extracellular contents (Fig 4C).

Prophenoloxidase (PO) activating system is an important identification and defense system for eliminating viruses in crustaceans (21). To find out whether Caspase 1 mediated cell death could influence PO system, Caspase 1 and IL-1β like protein were respectively knocked down and then the PO activity was measured with or without WSSV challenge. As expected, WSSV challenge greatly induced the PO activity (Fig 4D). However, either Caspase 1 knockdown or IL-1β like protein knockdown partially inhibited the PO activation (Fig 4D), suggesting that Caspase 1 mediated cell death could enhance PO activity upon WSSV infection.

Taken the above data together, these findings demonstrate that the releasing of IL-1β like protein is dependent on the enzymatic activity of Caspase 1, resulting in the activation of PO system and cell damage which prevent shrimp from virus infection.

### Negative correlation of Caspase 1 and IL-1β like protein expressions with virus infection

To investigate the influence of Caspase 1-mediated cell death on virus infection, WSSV copy number was detected upon knockdown of either *Caspase 1* or *IL-1β like protein* gene in shrimp. The results showed that the silencing of Caspase 1 or IL-1β like protein significantly increased the copy number of WSSV compared to the control, indicating that Caspase 1-mediated cell death played an important antiviral role in shrimp immunity (Fig 5A), which was further demonstrated by the shrimp cumulative mortality analysis (Fig 5B).

Collectively, these findings reveal a novel pathway in which Caspase 1-mediated cell death induces PO activation and plays a critical antiviral role in shrimp immunity (Fig 5C).

**Fig 5.**
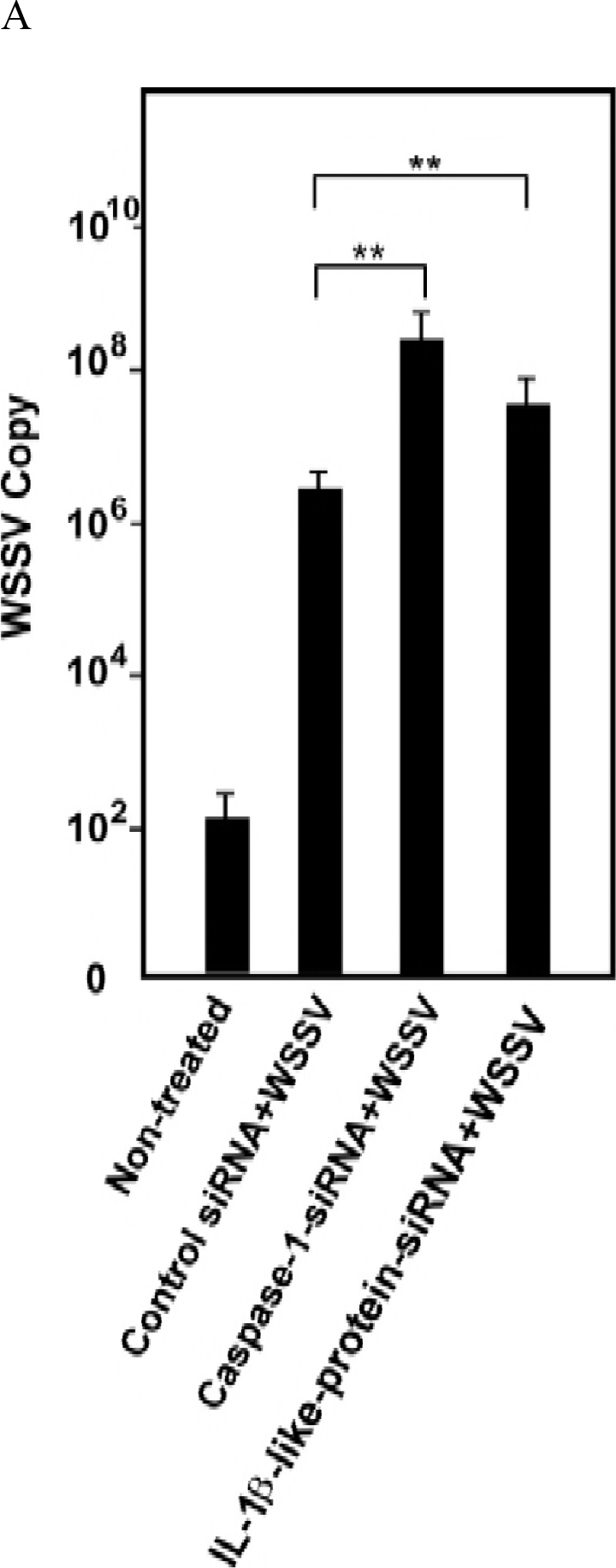
Negative correlation of Caspase 1 and IL-1β like protein expressions with virus infection. (A) Detection of WSSV copies after knockdown of Caspase 1 or IL-1β like protein genes. WSSV copy number in Caspase-1-siRNA or IL-1β-like-protein-siRNA treated shrimp was measured by quantitative real-time PCR. (B) Monitoring of shrimp cumulative mortality upon knockdown of Caspase 1 or IL-1β like protein genes. (C) A model for the Caspase 1-mediated antiviral pathway in shrimp.

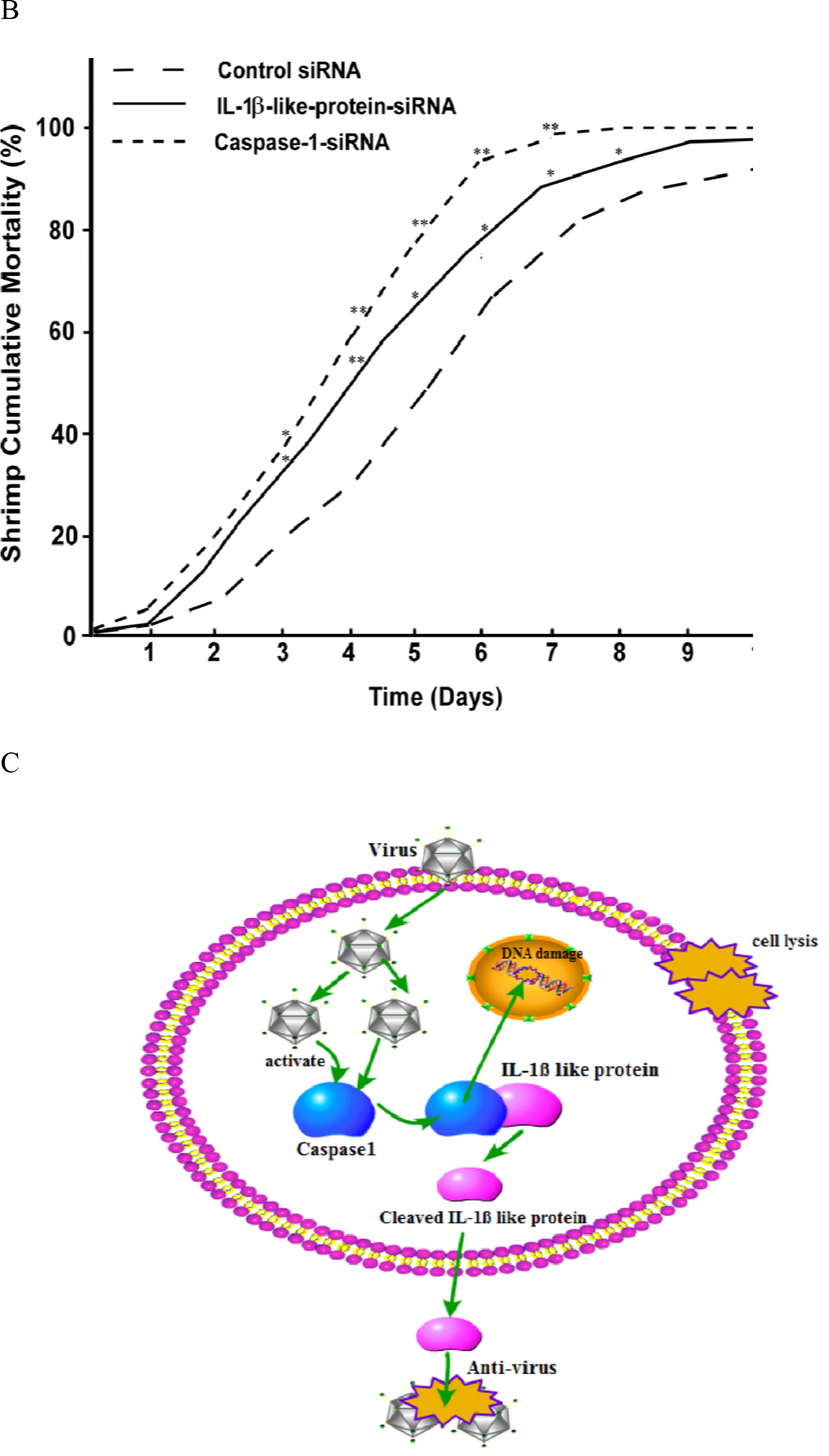

## Discussion

The outbreak of viral diseases is a major concern preventing the development of shrimp aquaculture industry. WSSV is one of the most virulent shrimp viruses and the processes of pathogenesis are extremely complex. Due to the lack of effective therapeutics to control virus, it is of great importance to understand viral pathogenesis and host responses at the molecular level in order to prevent virus invasion. So far, some studies have demonstrated that many genes and pathways are involved in host defense against WSSV, including Ran protein, the Ras-activated endocytosis process, the RNA interference pathway and apoptosis (22–25). Apoptosis is maybe the best described form of cell death that is mediated by the activation of the apoptotic caspase enzymes Caspase 3/7 and the antiviral role of apoptosis has been discovered both in invertebrates and vertebrates. Currently, pyroptosis, which is mediated by Caspase 1 activation, is thought to have the capacity of eliminating viruses only in vertebrates (26). In vertebrates, pyroptosis activation can result in not only the production of activated inflammatory cytokines, but also rapid cell death characterized by plasma-membrane rupture and release of proinflammatory intracellular contents(26). On the other hand, viruses including HIV have evolved mechanisms to inhibit pyroptosis, enhancing their ability to persist and cause disease (27). Ultimately, it is the competition between host and pathogen to regulate pyroptosis, and the outcome dictates life or death to the host or to viruses. Based on the critical role of Caspase 1 in the defense of a wide range of virus infections in vertebrates, in this study, Caspase 1 gene was first cloned from shrimp and the activity of Caspase 1 was then characterized. Our data showed that Caspase 1 activity was upregulated upon WSSV infection and the knockdown of Caspase 1 facilitated virus infection. Compared with the function of pyroptosis in vertebrates, our study revealed that Caspase 1-mediated cell death also contributed to eliminating virus in crustacean. Our findings indicated that the antiviral function of Caspase 1 was conservative although the sophisticated immune responses of shrimp have not been developed yet.

In vertebrates, the activated Caspase 1 recognizes and cleaves IL-1β into the mature form. The release of mature IL-1β further enhances the inflammatory response by stimulating immune cell activation and more cytokine secretion (28), leading to critical physiological consequences such as fever and hypotension. However, the cleavage and release of IL-1β is not indispensible for caspase 1-mediated cell death as demonstrated by IL-1β-/- knockout animals (29). In our study, the substrate of Caspase 1, which was named IL-1β like protein, was firstly identified. The results showed that WSSV infection induced the redistribution of IL-1β like protein into plasma of shrimp, indicating that IL-1β like protein was released into extracellular contents upon virus infection. The released IL-1β like protein, instead of triggering inflammation responses, activated phenoloxidase system of shrimp, an important invertebrate defense mechanism against virus infection. These data demonstrated that the function IL-1β like protein participating in Caspase 1-mediated cell death in shrimp was similar with that in vertebrates, and that IL-1β like protein functioned as cytokine-like molecules in invertebrates. Although invertebrates including shrimp lack adaptive immune system, several putative functional analogues of inflammatory cytokines have been identified so far. For example, tumor necrosis factor (TNF)-like molecules and interleukin-1 (IL-1)-like molecules are detected in a variety of invertebrates, such as insects, echinoderms and protochordates (30, 31). Moreover, chemokine IL-8 and transforming growth factor β1 are vertebrate cytokines involved in inflammatory processes and molecules crossreacting with the two proteins using antibodies have been identified in mollusks (32). In addition, IL-2-like activity is detected in protochordates and echinoderms which have hematopoietic organs and T-like cells (33). In this study, our data suggested that an invertebrate cytokine network might be operative and regulate host defense mechanisms against virus infection as in vertebrates.

## Material and methods

### Shrimp culture and WSSV challenge

*Marsupenaeus japonicas* shrimp was cultured in groups of 20 individuals in tanks containing aerated seawater at room temperature. The individual shrimp was about 10g in weight and 10 to 12 cm in length. To ensure the absence of WSSV in shrimp prior to experimental infection, PCR was performed using WSSV specific primers (5’-TATTGTCTCTCCTGACGTAC-3’ and 5’-CA CATTCTTCACGAGTCTAC-3’). Virus-free shrimp were infected with WSSV (10^5^ copies/ml) by injection into the lateral area of the fourth abdominal segment. Three shrimp were randomly collected for each treatment. At different time post-infection, the shrimp hemolymph was collected for later use.

### Rapid amplification of cDNA ends (RACEs)

Total RNAs were extracted from shrimp hemocytes using mirVana miRNA™ Isolation Kit (Ambion, USA). RACE experiments were conducted using 5’-3’ RACE Kit (Roche, USA) according to the manufacturer’s protocol. Briefly, cDNA was synthesized and applied for PCR and nest-PCR. PCR was performed using 3’ RACE primer, 5’ RACE primer or/and gene-specific primers. The amplified DNAs were sequenced.

### Quantitative real-time PCR for detections of virus copies and gene expression

Quantitative real-time PCR was performed to quantify WSSV copies and gene expression level in shrimp. DNA or RNA was extracted from shrimp hemocytes using SQ tissue DNA kit (Omega-BioTek, USA) or RNA Isolation kit (Ambion, USA). RNA was reversely transcribed into cDNA by cDNA synthesis kit (Takara, Japan).

The WSSV copies were evaluated with WSSV-specific primers (5’-TTGGTTTCAG CCCGAGATT-3’ and 5’-CCTTGGTCAGCCCCTTGA-3’). The gene expression level was determined using gene-specific primers (Caspase 1, 5’-TTTCAAGGCTGACCAGT GCCCTAC-3’ and 5’-ACCTGCGTCAAGACCCTCACCTC-3’; IL-1β, 5’-GCGTTCCACATTTCAGATACTCGG-3’ and 5’-TGTAGACCATTCCAAAGGCTCCC-3’). The 25μL PCR solution contained 12.5μL of 2×Premix Ex Taq (TaKaRa, Japan), 0.5μL of 10μM forward primer, 0.5μL of 10μM reverse primer and 1μM DNA template. The stages of PCR program were 95°C for 5 min, followed by the amplification stage consisting of 40 cycles of 95°C for 10 s and 60°C for 30 s.

### Western blot analysis

Proteins were separated by a 12% SDS-polyacrylamide gel, which were then transferred onto a polyvinylidene difluoride (PVDF) membrane (Millipore, USA). The membrane was blocked with 5% nonfat milk in phosphate buffered solution (PBS) for 2 h at room temperature. The blot was incubated with a primary antibody at 4°C overnight. After washes with PBS, the blot was incubated with the fluorescence-labeled anti-mouse IgG (Cell Signaling Technology, USA) for 2 h at 4°C. The blot was detected by Odyssey (Licor, USA) for signals.

### Caspase 1 activity detection

Caspase 1 Activity Assay Kit was used to detect Caspase 1 activity according to the manufacturer’s instructions (Beyotime, China). Briefly, shrimp hemocytes were collected and washed with PBS. Then, the hemocytes were lysed in 100μL ice-cold lysis buffer for 15 min and centrifuged at 15,000×g (4°C) for 10 min. The supernatant was incubated with the substrate Ac-YVAD-pNA (2mM) for 2 h at 37°C. The reaction was detected by a spectrophotometer at OD_405_. Caspase 1 inhibitor Belnacasan was used to inhibit Caspase 1 activity in shrimp. The inhibitor was injected into shrimp at 100μg/g shrimp in combination with WSSV injection or not.

### Cell culture and plasmids transfection

Insect High Five cells (Invitrogen, USA) were cultured and maintained at 27°C in Express Five serum-free medium (Invitrogen) containing L-glutamine (Invitrogen). Cells were cotransfected with plasmids expressing Flag-Caspase 1 and His-IL-1β like protein according to the manufacture’s instruction. Briefly, when the cells reached about 70% confluence in a 6-well plate, 2μg for each indicated plasmid and 6μl Cellfectin Reagent were diluted in 100 μl Grace’s medium, respectively. The diluted plasmids and Cellfectin Reagent were then mixed gently and incubated for 30 minutes at room temperature. The mixture was added to the cells and incubated in a 27°C incubator for 5 h before the medium was replaced by fresh Express Five serum-free medium. Cells were finally subjected to further research 48 h after transfection.

### Co-immunoprecipitation

Cells were collected and lyzed in ice-cold lysis buffer (50mM Tris-HCl, 150mM NaCl, 0.2% NP40, 1mM PMSF, 1×protease inhibitor, pH 8.0). The lysate was then centrifuged three times for 5 min each to remove all the undissolved contents. The supernatant was incubated with indicated antibody overnight at 4°C. Protein A+G agarose beads (Invitrogen, USA) were incubated with the lysate for 2 h at 4°C. After washed three times with ice-cold wash buffer (50mM Tris-HCl, 200mM NaCl, 0.1% NP40, 1mM PMSF, 1×protease inhibitor, pH 8.0), the immuno-complexes were eluted by 100 mM glycine and subjected to Western blotting for the detection of indicated proteins.

### Synthesis of siRNAs and RNAi assay

Based on the sequences of *Caspase 1* and *IL-1β like protein* genes, siRNAs were designed to specifically target the genes and then were synthesized in vitro using a commercial kit according to the manufacturer’s instructions (TaKaRa, Japan). The siRNAs used were Caspase 1-siRNA (5’-AAACCACUCGGAAUGCGAUGCG-3’) and IL-1β-like-protein-siRNA (5’-GGGAAGGCTTTGAGGTCCTTGT-3’). The scrambled siRNA not targeting any genes was used as the control. The formation of double-stranded RNAs was monitored by determining the size in agarose gel electrophoresis. The synthesized siRNAs were dissolved in siRNA buffer (50 mM Tris-HCl, 100 mM NaCl, pH 7.5) and quantified by Nanodrop 2000. The RNA interference (RNAi) assay was conducted in shrimp by the injection of a siRNA into the lateral area of the fourth abdominal segment at 30 μg/shrimp using a syringe with a 29-gauge needle. The siRNA (15μg) and WSSV (10^5^ copies/ml) were co-injected into virus-free shrimp at a volume of 100μL per shrimp. At 12 h after the co-injection, the siRNA (15μg) (100μL /shrimp) was injected into the same shrimp. For each treatment, 20 shrimp were used. At different times after the last injection, the shrimp hemocytes were collected. Three shrimp specimens from each treatment, randomly selected, were collected for analysis. The assays were biologically repeated for three times.

### Phenoloxidase activity detection

Hemolymph was centrifuged and the plasma was used for the detection of phenoloxidase activity. The reaction buffer (0.1M K3PO4, 0.01M L-dopa, pH 6.0) and plasma was mixed and then incubated at 28°C for 2 min. The activity of phenoloxidase was measured by a spectrophotometer at OD492.

### Shrimp mortality analysis

Shrimp with different treatments were cultured at 20 shrimp/treatment. Shrimp mortality was examined every day. The experiment was biologically repeated three times.

### Statistical analysis

The data from three independent experiments were analyzed by one-way analysis of variance (ANOVA) to calculate the mean and standard deviation (SD) of the triplicate assays.

